# Variation at the *TRIM11* locus modifies Progressive Supranuclear Palsy phenotype

**DOI:** 10.1101/333195

**Authors:** E Jabbari, J Woodside, MMX Tan, M Shoai, A Pittman, R Ferrari, KY Mok, D Zhang, RH Reynolds, R de Silva, MJ Grimm, G Respondek, U Müller, S Al-Sarraj, SM Gentleman, AJ Lees, TT Warner, J Hardy, T Revesz, PROSPECT-UK consortium, GU Höglinger, JL Holton, M Ryten, HR Morris, PROSPECT-UK consortium

## Abstract

**Objective:** The basis for clinical variation related to underlying Progressive Supranuclear Palsy (PSP) pathology is unknown. We performed a genome wide association study (GWAS) to identify genetic determinants of PSP phenotype.

**Methods:** Two independent pathological and clinically diagnosed PSP cohorts were genotyped and phenotyped to create Richardson’s syndrome (RS) and non-RS groups. We carried out separate logistic regression GWAS to compare RS and non-RS groups and then combined datasets to carry out a whole cohort analysis (RS=367, non-RS=130). We validated our findings in a third cohort by referring to data from 100 deeply phenotyped cases from a recent GWAS. We assessed the expression/co-expression patterns of our identified genes and used our data to carry out gene-based association testing.

**Results:** Our lead single nucleotide polymorphism (SNP), rs564309, showed an association signal in both cohorts, reaching genome wide significance in our whole cohort analysis – OR 5.5 (3.2-10.0), p-value 1.7×10^−9^. rs564309 is an intronic variant of the tripartite motif-containing protein 11 (*TRIM11*) gene, a component of the ubiquitin proteasome system (UPS). In our third cohort, minor allele frequencies of surrogate SNPs in high linkage disequilibrium with rs564309 replicated our findings. Gene based association testing confirmed an association signal at *TRIM11*. We found that TRIM11 is predominantly expressed neuronally, in the cerebellum and basal ganglia.

**Interpretation:** Our study suggests that the *TRIM11* locus is a genetic modifier of PSP phenotype and potentially adds further evidence for the UPS having a key role in tau pathology, therefore representing a target for disease modifying therapies.

## Introduction

Progressive supranuclear palsy (PSP) is a progressive neurodegenerative condition and the most common cause of atypical parkinsonism, with an estimated prevalence of 5-7 per 100,000 (1). The pathology of PSP is centred on the structural microtubule associated protein tau, encoded by the *MAPT* gene located on chromosome 17. In PSP there is neuronal and glial accumulation of hyper-phosphorylated fibrillary aggregates of 4-repeat (4R) predominant tau. The pathological hallmarks of PSP include a high density of neurofibrillary tangles and neuropil threads in the basal ganglia and brainstem along with tau-positive tufted astrocytes (2).

Richardson’s syndrome (RS) is the most common clinical phenotype related to PSP pathology – first described by Steele, Richardson and Olszewski as a levodopa-unresponsive akinetic-rigid syndrome with falls, a vertical supranuclear gaze palsy and dementia (3). Previous studies looking at the natural history of RS have shown that the mean age of disease onset is 65-67 years, and the median disease duration is 6-7 years (4). In addition, a clinical diagnosis of RS has been shown to be highly predictive of underlying PSP pathology (5), and the diagnosis of this form of PSP was operationalised in the NINDS-SPSP criteria (6).

We and others have identified alternative clinical phenotypes (7, 8) related to PSP-pathology in relatively small case series. This led to the description of two distinct PSP non-RS clinical phenotypes by Williams and colleagues, PSP-Parkinsonism (PSP-P) (9) and Pure Akinesia with Gait Freezing (PAGF) (10). PSP-P and PAGF have a similar age of disease onset to RS, clinically resemble RS in the latter stages of disease and have a significantly longer mean disease duration (PSP-P = 9 years; PAGF = 13 years). The basis for this clinical variation related to a core pathology is unknown. PSP clinical subtypes have been related to the regional distribution and severity of pathogenic tau accumulation and neuronal loss (11). Although post-mortem remains the gold standard for diagnosing PSP, recent publication of new diagnostic criteria from the Movement Disorder Society (MDS) PSP study group (12) highlight the presence of PSP-P and PAGF along with other PSP clinical phenotypes relating to underlying PSP pathology including PSP-corticobasal (PSP-CBS) (13) and PSP-frontal (PSP-F) subtypes (14).

A recent comprehensive genome wide association study (GWAS) involving 1,114 pathologically confirmed PSP cases and 3,247 controls was carried out to identify common risk variants for PSP. Single nucleotide polymorphisms (SNPs) that passed a significance cut off point of p≤10^−3^ were subsequently genotyped in a validation cohort that consisted of 1,051 clinically diagnosed PSP cases and 3,560 controls. Loci at MAPT (H1 haplotype and H1c sub-haplotype), MOBP, STX6 and EIF2AK3 were associated with PSP (15).

Differences in the clinico-pathological phenotypes of tauopathies (including Alzheimer’s disease) may relate to differences in the strain properties of toxic tau species (16).

However, here we use a large clinico-pathological cohort based on the latest MDS diagnostic criteria to show that the clinical phenotype of PSP relates in part to genetic variants which may determine regional susceptibility.

## Methods

### Study design and participants

All patients gave written informed consent for the use of their medical records and blood/brain tissue for research purposes, including the analysis of DNA. Patients with a neuropathological diagnosis of PSP were identified from the following UK brain banks: MRC London Neurodegenerative Diseases brain bank (Research Ethics Committee reference 08/MRE09/38+5); Multiple Sclerosis and Parkinson’s brain bank, London (Research Ethics Committee reference 08/MRE09/31+5); Queen Square brain bank (the brain donor programme was approved by a London Multi-Centre Research Ethics Committee and tissue is stored for research under a license from the Human Tissue Authority, No. 12198). The year of death for cases ranged from 1998-2017.

Patients with a clinical diagnosis of a PSP syndrome were identified from the Progressive Supranuclear Palsy Cortico-Basal Syndrome Multiple System Atrophy Longitudinal UK (*PROSPECT*-UK) study, a longitudinal study of patients with atypical parkinsonian syndromes undergoing serial clinical, imaging and biomarker measures: Queen Square Research Ethics Committee 14/LO/1575. Cases were recruited between 2015 and 2017. A subset of these patients also underwent post-mortem neuropathological diagnosis at the Queen Square brain bank.

### Phenotyping of cases

Retrospective clinical notes review of all neuropathological PSP cases was performed in order to extract the following demographic and clinical information: gender; age at motor symptom onset; date of motor symptom onset; date of death. This information was used to calculate the total disease duration (defined as: date of death – date of motor symptom onset). Cases that did not have the above clinical information available were excluded from the study. Exclusion criteria used in the MDS diagnostic criteria were not considered as the presence of alternative diseases would have been identified at post-mortem. Using the MDS diagnostic criteria, each case was assigned an initial and final clinical phenotype (12). This was based on the clinical features documented in clinical letters in the first 3 years from motor symptom onset and the clinical features documented in clinical letters in the last 2 years of life. We focused on three clinical phenotypes of interest: RS, PSP-P and PAGF; and only assigned these phenotypes if their corresponding “probable” criteria were fulfilled. Cases were assigned a diagnosis of ‘unclassified’ if there was insufficient evidence from the clinical notes to assign one of the phenotypes of interest. In cases where there was an overlap of clinical phenotype features, a consensus decision was made to assign the most appropriate clinical phenotype. The same clinical data, as above, was collected on clinically diagnosed PSP cases using their PROSPECT-UK study clinical assessments. To ensure accuracy in assigning a phenotype, living subjects were only included if their latest clinical assessment was carried out at least 3 years after motor symptom onset. In addition, cases were excluded from analyses if they had the presence of any MDS diagnostic exclusion criteria or if they fulfilled both MDS criteria for one of our PSP phenotypes of interest as well as Armstrong criteria for probable CBS as these subjects may have underlying Corticobasal Degeneration (CBD) pathology (17).

### Genotyping and quality control

All pathologically diagnosed cases had DNA extracted from frozen brain tissue (cerebellum or frontal cortex). Subsequently, DNA samples from all cases underwent genotyping using the Illumina NeuroChip (18). Standard genotype data quality control steps were carried out as per Reed et al (19), including a principal components analysis (PCA) to exclude all non-European subjects. All cases were screened for known *MAPT, LRRK2 and DCTN1* mutations covered by the NeuroChip. SNP imputation was carried out on our NeuroChip data using the Sanger Imputation Service to produce a final list of common (minor allele frequency ≥1%) variants for analyses. Imputed SNP positions were based on Genome Reference Consortium Human 37/human genome version 19 (GRCh37/hg19). Standard quality control steps taken for SNP imputation were carried out as per Reed et al (19).

To confirm the validity of our NeuroChip genotyping and imputation, a subset of both directly genotyped and imputed SNPs underwent re-genotyping using the LGC KASP genotyping service for coverage of significant regions in association.

### Statistical analyses

All statistical analyses were carried out using Plink version 1.9 and images generated using R version 3.3.2 and LocusZoom.

By dividing the whole cohort into RS and non-RS (combined PSP-P and PAGF) groups based on their initial clinical phenotype, group comparisons of clinical features were carried out using t-tests. In addition, the RS and non-RS group minor allele frequencies (MAF) of all PSP case-control GWAS risk variants were extracted from our imputed data.

### Logistic regression GWAS

A logistic regression GWAS was performed on our imputed data to compare RS and non-RS groups. Based on their assigned initial clinical phenotypes, non-RS subjects were defined as “cases” and RS subjects were defined as “controls.” The regression model used gender, age at motor symptom onset, study site of subject recruitment and the first two principal components as covariates. This analysis was first carried out on our pathological cohort and then subsequently on our clinical cohort before combining datasets to carry out a whole cohort analysis. The Bonferroni correction for multiple SNP testing was used to set the genome wide significance p-value threshold at 9×10^−9^. The whole cohort GWAS analysis was used to generate Manhattan and regional association plots.

All significant SNPs from our association analysis were assessed for their MAFs in European controls. This data was acquired from the Genome Aggregation Database (www.gnomad.broadinstitute.org) which is based on data from ~120,000 exome sequences and ~15,500 whole-genome sequences from unrelated individuals.

All significant SNPs from our association analysis were assessed for their level of significance in phase 1 of the original PSP case-control GWAS (15) using publically available data at The National Institute on Aging Genetics of Alzheimer’s Disease Data Storage Site (www.niagads.org).

### PSP case-control GWAS validation cohort

A separate subset of 100 pathologically confirmed PSP cases from phase 1 of the original PSP case-control GWAS had in-depth phenotype data available to assign an initial clinical phenotype according to the MDS criteria, as per our study methods. These cases had undergone genotyping using the Illumina Human 660W-Quad Infinium Beadchip with standard data quality control steps taken, including a PCA to exclude non-Europeans. RS and non-RS group MAFs for directly genotyped SNPs that were significant in our phenotype GWAS were extracted to further validate our findings.

#### Gene-based association testing

Gene-level p-values were calculated using MAGMA v1.06 as outlined in de Leeuw et al (20). MAGMA tests the joint association of all SNPs in a gene with the phenotype while accounting for linkage disequilibrium between SNPs. This presents a powerful alternative to SNP-based analyses, as it reduces the multiple testing burden and thus increases the possibility of detecting effects consisting of multiple weaker associations (20). SNPs were mapped to genes using NCBI definitions (GRCh37/hg19, annotation release 105); only genes in which at least one SNP mapped were included in downstream analyses. These were run both with and without a window of 35 kb upstream and 10 kb downstream of each gene, as most transcriptional regulatory elements fall within this interval (21). Furthermore, the MHC region was excluded. The gene p-value was computed based on the mean association statistic of SNPs within a gene, with genome wide significance set to p-value < 2.74×10^−6^, and linkage disequilibrium was estimated from the European subset of 1000 Genomes Phase 3.

#### Whole exome sequencing

69 cases from our pathological cohort had previously undergone whole exome sequencing (WES) using the Illumina Truseq Capture in Illumina HiSeq platform. This data was used to look for the presence of rare coding variants in genes of interest to arise from our GWAS. Read data was aligned to hg19 by use of novoalign (V3.02.04) and indexed bam files were de-duplicated of PCR artefacts by use of Picard Tools MarkDuplicates. The Genome Analysis Toolkit (GATK) was then used to perform all subsequent steps according to their good practice; local realignments around possible indels, variant calling was conducted with HaplotypeCaller. Variants were filtered by use of variant quality score recalibration (truth tranche 99.0%). In addition, hard-filtering based on low-depth and low-genotype quality was performed. Annotation was performed by use of Annovar software.

#### Assessment of gene expression

Gene expression profiles were assessed using publically available BRAINEAC (www.braineac.org) (22), GTEx (www.gtexportal.org) and Allen Brain Atlas (www.brain-map.org) (23) web based resources.

The BRAINEAC database contains brain tissues from 134 healthy controls from the following brain regions: frontal cortex, temporal cortex, parietal cortex, occipital cortex, hippocampus, thalamus, putamen, substantia nigra, medulla, cerebellum and white matter. RNA isolation and processing of brain samples was performed and analysed using Affymetrix Exon 1.0 ST arrays. The GTEx database consists of 8555 samples from 53 tissues (including 13 brain regions) of 544 donors for which RNAseq was conducted. The GTEx Project was supported by the Common Fund of the Office of the Director of the National Institutes of Health, and by NCI, NHGRI, NHLBI, NIDA, NIMH, and NINDS. The data used for the analyses described in this manuscript were obtained from the GTEx Portal on 04/31/18. The Allen Human Brain Atlas database contains microarray data from 8 neuropathologically normal individuals of varying ethnicity. Microarray data was generated using the Agilent 4×44 Whole Human Genome array and covers ~62,000 gene probes per profile and ~150 brain regions.

Gene expression at the cellular level in the brain was analysed using RNAseq data from the Brain RNA-Seq database (www.brainrnaseq.org/) as per Zhang et al (24). Of note, this data was generated from healthy temporal lobe samples that were resected from 14 patients in order to gain access to deeper epileptic hippocampi. The number of different cell types obtained from these samples were as follows: mature astrocyte, n = 12; microglia, n = 3; neuron, n = 1; oligodendrocyte, n =3. We also used single cell RNA-seq data provided by DropViz (www.dropviz.org), which provides gene expression on 690,000 individual cells derived from nine different regions of the adult mouse brain (25).

#### Colocalisation analyses

To evaluate the probability that the same causal SNP was responsible for modifying the phenotype of PSP and modulating gene expression, we performed the Coloc method described by Giambartolomei et al (26) using our GWAS summary statistics coupled with eQTLs from Braineac and GTEx. GTEx eQTLs included those originating from all brain regions. We restricted analyses to genes within 1Mb of the significant region of interest (p-value < 5×10^−8^) and ran coloc.abf with default priors. We considered tests with a PPH4 >= 0.75 to have strong evidence for colocalisation.

### Results

A total of 497 subjects were included for analyses. Their clinical features are summarised by cohort and disease group in the table below (Table 1).

**Table 1:** Clinical features of subjects included in genotype-phenotype analyses. RS/non-RS status based on initial clinical phenotype. No statistically significant difference between RS and non-RS groups. No statistically significant difference between pathological and clinical cohorts. † Statistically significant (p < 0.05) difference between RS and non-RS groups using Welch’s t-test.

|  | <u>Pathological cohort</u> |  | <u>Clinical cohort</u> |  | <u>Whole cohort</u> |  |
| --- | --- | --- | --- | --- | --- | --- |
|  | RS | non-RS | RS | non-RS | RS | non-RS |
| No. of subjects | 230 | 76<br>PSP-P = 60<br>PAGF = 16 | 137 | 54<br>PSP-P = 42<br>PAGF = 12 | 367 | 130<br>PSP-P = 102<br>PAGF = 28 |
| % of subjects male | 60.0% | 53.9% | 57.7% | 51.9% | 59.1% | 53.1% |
| Age at motor symptom onset (years)–<br>Mean, range, SD | 68.9<br>49-89<br>7.4 | 65.9<br>46-86<br>8.8 | 66.5<br>51-87<br>6.8 | 65.3<br>54-82<br>6.9 | 68.1 □<br>□<br>49-89<br>7.3 | 65.6 □ □<br>46-86<br>8.0 |
| Final/current clinical phenotype | RS =<br>230 | RS = 71<br>PSP-P = 4<br>PAGF = 1 | RS =<br>137 | RS = 28<br>PSP-P = 20<br>PAGF = 6 | RS =<br>367 | RS = 99<br>PSP-P = 24<br>PAGF = 7 |
| Mean disease duration in deceased<br>subjects (years)–Mean, range, SD | 5.9<br>1.9-15.9<br>1.9 | 10.7<br>2.2-16.3<br>2.9 | 5.6<br>2.4-13.5<br>2.2 | 9.2<br>8.1-10.8<br>1.2 | 5.8 ⊥ †<br>1.9-15.9<br>1.9 | 10.6 ⊥ †<br>2.4-13.5<br>2.8 |
| No. of subjects undergoing post-mortem<br>(% with a pathological diagnosis of PSP) | 230<br>(100%) | 76<br>(100%) | 10<br>(100%) | 1<br>(100%) | 240<br>(100%) | 77<br>(100%) |

44 subjects were deemed unclassifiable and therefore not included in subsequent analyses. An initial screen of our genotype data revealed similar MAFs for risk variants identified in the PSP case-control GWAS (Table 2).

**Table 2:** Comparison of PSP risk variant status between PSP case-control GWAS and PSP phenotype GWAS data. PSP case-control GWAS data taken from Höglinger et al (16). † No statistically significant difference between RS and non-RS groups using Fisher’s exact test.

| Chr.<br>band | SNP<br>Position<br>(BP) | Gene | PSP case-control GWAS |  | PSP phenotype GWAS |  |
| --- | --- | --- | --- | --- | --- | --- |
|  |  |  | MAF in healthy<br>controls | MAF in<br>PSP | MAF in<br>RS | MAF in non-RS |
| 1q25.3 | rs1411478<br>180,962,282 | *STX6* | 0.42 | 0.50 | 0.44† | 0.43 |
| 2p11.2 | rs7571971<br>88,895,351 | *EIF2AK3* | 0.26 | 0.31 | 0.34† | 0.30 |
| 3p22.1 | rs1768208<br>39,523,003 | *MOBP* | 0.29 | 0.36 | 0.32† | 0.32 |
| 17q21.31 | rs8070723<br>44,081,064 | *MAPT* (H1<br>haplotype) | 0.23 | 0.05 | 0.05† | 0.05 |
|  | rs242557<br>44,019,712 | *MAPT*<br>(H1c sub-haplotype) | 0.35 | 0.53 | 0.44† | 0.49 |

In addition, none of our cases carried known pathogenic variants covered by the NeuroChip for the following genes: *MAPT*(40 variants); *LRRK2* G2019S (1 variant); *DCTN1* (12 variants).

After SNP imputation and data quality control, 6,215,948 common variants were included in our analysis. By assigning non-RS subjects as “cases” and RS subjects as “controls” we applied a logistic regression association analysis using gender, age at motor symptom onset, study site of subject recruitment and the first two principal components as covariates. We first carried out this analysis using data from our pathological cohort and then validated our findings using data from our independent clinical cohort before combining both datasets for a whole cohort analysis. The whole cohort analysis revealed 27 SNPs, all located on chromosome 1, which passed the threshold for genome-wide significance (p-value < 9×10^−9^). These results are summarised in Figure 1. Population stratification was not evident in our cohort as non-European subjects were excluded from analyses as part of our genotype data quality control. This was further confirmed by obtaining a genomic inflation factor (lambda) value of 1.05. A further locus on chromosome 12 approached genome wide significance with the lead SNP (rs621042) p-value at 7.8×10^−7^.

**Figure 1:**
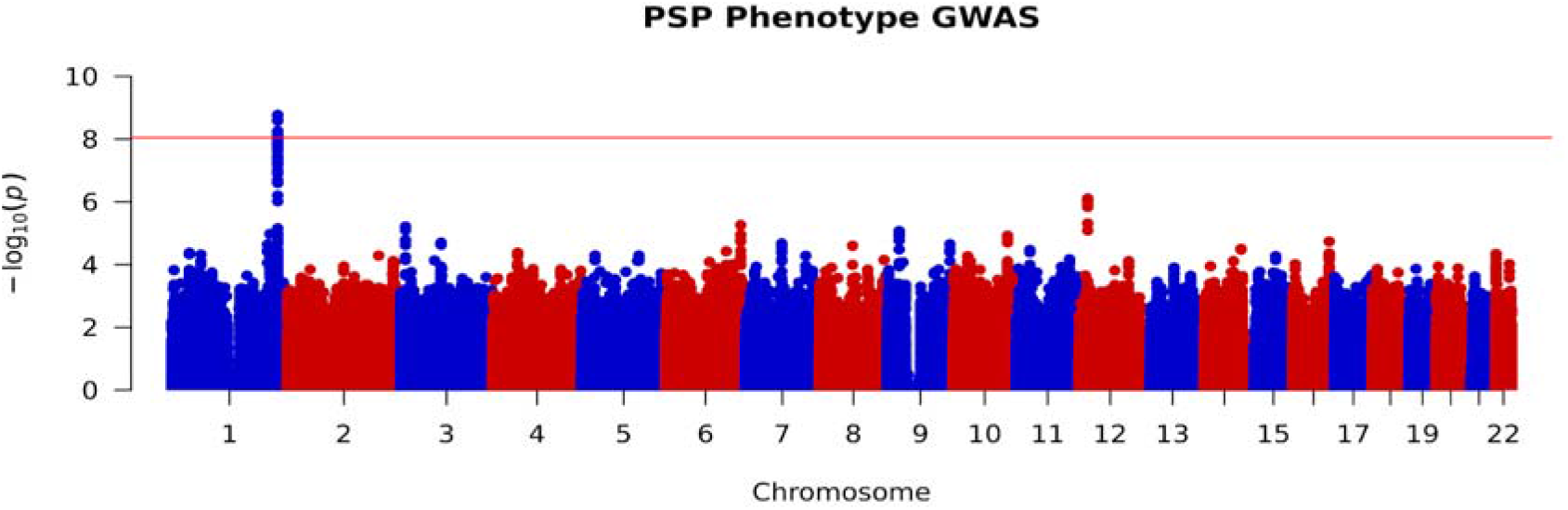
Manhattan plot of whole cohort RS vs non-RS association analysis, highlighting genome-wide significance at chromosome 1. Red line indicates threshold for genome wide significance (p-value < 9×10^−9^).

An in-depth analysis of our significant SNPs reveal that they are all in high LD with each other (as defined by r^2^ > 0.80) and are all located at the chromosome 1q42.13 locus. A regional association plot (Figure 2) reveals that our lead SNP, rs564309, is an intronic variant located in between exons 3 and 4 of the tripartite motif-containing protein 11 (*TRIM11*) gene. Alongside our directly genotyped lead SNP, the imputation quality (INFO) score for imputed significant SNPs ranged from 0.96 to 1. 96 cases from our pathological cohort underwent re-genotyping for 8 SNPs (rs564309, rs35670307, rs12065815, rs10158354, rs3795811, rs6426503, rs138782220 and rs7555298) that span the significant chromosome 1q42.13 locus. 3 of the 8 SNPs, including our lead SNP, were originally directly genotyped via the NeuroChip while the remaining 5 SNPs were originally imputed in our dataset. The p-value of these SNPs in our whole cohort GWAS ranged from 1.7×10^−9^ to 7.3×10^−5^. The results of this re-genotyping run showed 100% concordance with our original NeuroChip and imputation data.

**Figure 2:**
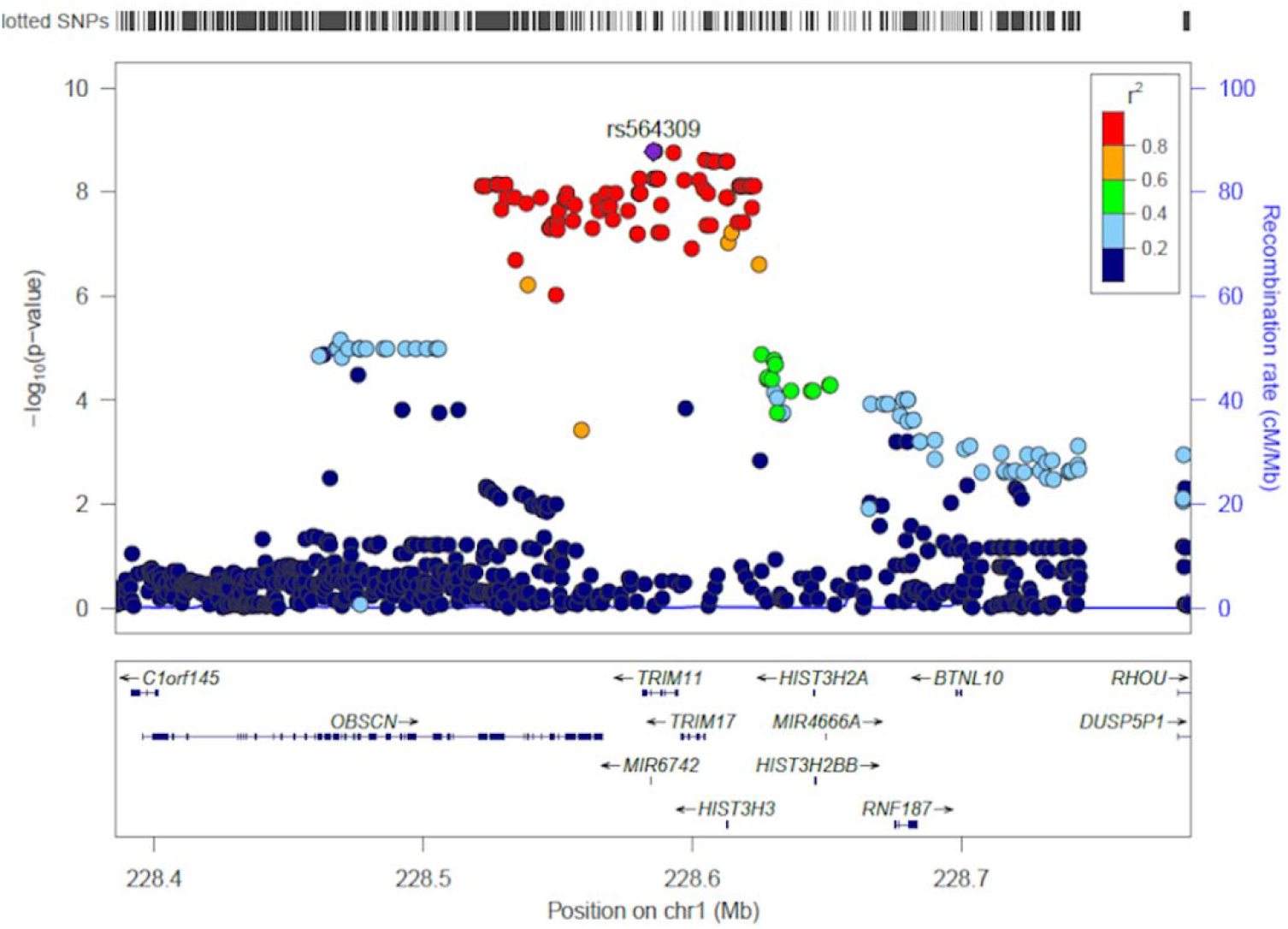
Regional association plot of RS vs non-RS association analysis using imputed SNP data, implicating the chromosome 1q42.13 locus and identifying rs564309, an intronic variant of *TRIM11*, as our lead SNP. SNP positions, recombination rates and gene boundaries based on GRCh37/hg19.

Furthermore, three of the significant SNPs in LD with rs564309 are non-synonymous (missense) coding variants of the Obscurin (*OBSCN*) gene. *OBSCN* is mainly expressed in skeletal muscle and may have a role in the organization of myofibrils during assembly as well as mediating interactions between the sarcoplasmic reticulum and myofibrils (27). Related diseases include fibromuscular dysplasia and hypertrophic obstructive cardiomyopathy.

The association statistics for rs564309, and the most significant flanking SNPs located at neighbouring genes within the chromosome 1q42.12 locus, are summarised below (Table 3). The MAF of these SNPs was shown to be 0.10 in healthy European controls on the gnomAD database.

**Table 3:** RS vs non-RS association statistics for rs564309, and the most significant flanking SNPs located at neighbouring genes, in pathological, clinical and whole cohorts respectively. SNP positions based on GRCh37/hg19. MAF = Minor Allele Frequency. OR = Odds Ratio. P = p-value in whole cohort analysis. RS = PSP-Richardson’s syndrome group. Non-RS = Combined PSP-Parkinsonism and Pure Akinesia with Gait Freezing group. † = p-value > 9×10^−9^.

| SNP<br>Position (BP) | Gene | Pathological cohort |  |  | Clinical cohort |  |  | Whole cohort |  |
| --- | --- | --- | --- | --- | --- | --- | --- | --- | --- |
|  |  | MAF<br>in RS | MAF in<br>non-RS | OR<br>(95% CI) | MAF<br>in RS | MAF in<br>non-RS | OR<br>(95% CI) | OR<br>(95% CI) | P |
| rs564309<br>228,585,562 | TRIM11 | 0.04 | 0.19 | 6.25†<br>(3.12-12.5) | 0.04 | 0.16 | 4.76†<br>(1.96-12.5) | 5.55<br>(3.22-10.0) | 1.7x10<sup>-9</sup> |
| rs61827276<br>228,597,130 | TRIM17 | 0.04 | 0.18 | 5.88†<br>(2.78-12.5) | 0.04 | 0.16 | 5.55†<br>(2.17-14.3) | 5.55<br>(3.12-10.0) | 6.2x10<sup>-9</sup> |
| rs61825312<br>228,530,748 | OBSCN | 0.04 | 0.19 | 5.88†<br>(2.86-12.5) | 0.04 | 0.16 | 4.35†<br>(1.78-11.1) | 5.26<br>(2.94-9.09) | 7.1x10<sup>-9</sup> |
| rs2230656<br>228,612,838 | HIST3H3 | 0.06 | 0.23 | 4.35†<br>(2.32 -8.33) | 0.06 | 0.20 | 3.70†<br>(1.75-8.33) | 4.00<br>(2.50-6.67) | 1.3x10<sup>-8</sup> |

To explore the impact of inadvertently including Parkinson’s disease (PD) cases in our clinically diagnosed non-RS group, we referred to genotyping data from 484 European clinically diagnosed PD cases that were genotyped alongside our PSP cases and had undergone the same quality control steps outlined above. We found that the MAF of rs564309 in PD cases was 7%, similar to the MAF in healthy controls and considerably lower than the MAF in our non-RS group.

When referring to publically available p-value data from phase 1 of the original PSP case-control GWAS, we found that none of our significant SNPs reached even nominal significance (p-value <0.05). 100 European pathologically confirmed PSP cases from this GWAS underwent retrospective phenotyping according to the MDS diagnostic criteria using available clinical notes. Of those, 83 cases fulfilled probable criteria for initial clinical phenotypes of relevance to this study (PSP-RS n = 45, PSP-P n = 38). rs1188473, a SNP that was directly genotyped in the case-control GWAS, in high LD with our lead SNP (r^2^ 1.0) and found to be significant in our phenotype GWAS (p-value 2.6×10^−9^), was shown to have similar MAFs when comparing the GWAS datasets in both RS (4% vs 6%) and non-RS (16% vs 16%) groups, therefore further validating our findings.

Analysis of WES data from 65 subjects (49 RS, 16 non-RS) within our pathological cohort did not identify any non-synonymous coding variants in *TRIM11* or *TRIM17* genes.

MAGMA analyses revealed that four genes passed genome-wide significance in analyses run with and without 35 kb upstream and 10 kb downstream of each gene (*TRIM11*, P = 5.64 × 10^−9^; *TRIM17*, P = 8.99 × 10^−9^; *HIST3H3*, P = 1.29 × 10^−8^, *LOC101927401*, P = 5.72 × 10^−8^) (Figure 3). *LOC101927401* appeared only in NCBI annotation and was absent in the queried gene expression databases, thus was excluded in downstream analyses.

**Figure 3:**
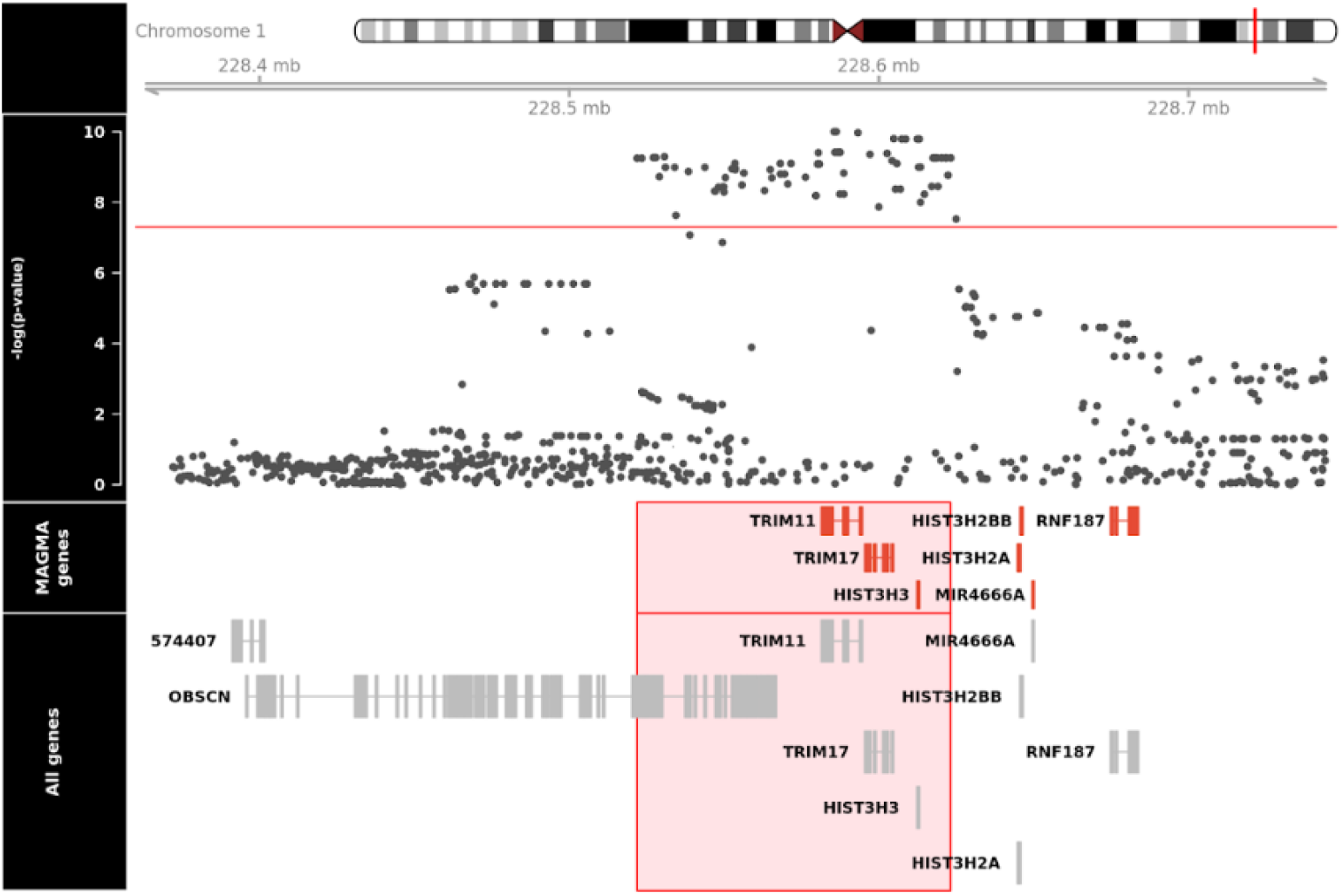
MAGMA analyses revealing significance at *TRIM11, TRIM17* and *HIST3H3* genes. Gene boundaries based on GRCh37/hg19. Red line indicates threshold for genome wide significance (p-value < 2.74×10^−6^).

We then used gene expression and co-expression data from BRAINEAC, GTEx and Allen Atlas databases to assess the expression profiles of genes identified in our MAGMA analysis: *TRIM11, TRIM17* and *HIST3H3*. All three datasets revealed high levels of TRIM11 and TRIM17 expression in the brain, particularly cerebellum and putamen, whereas HIST3H3 expression appeared to be at the lower limit of detection in human brain (Figure 4A-C).

**Figure 4:**
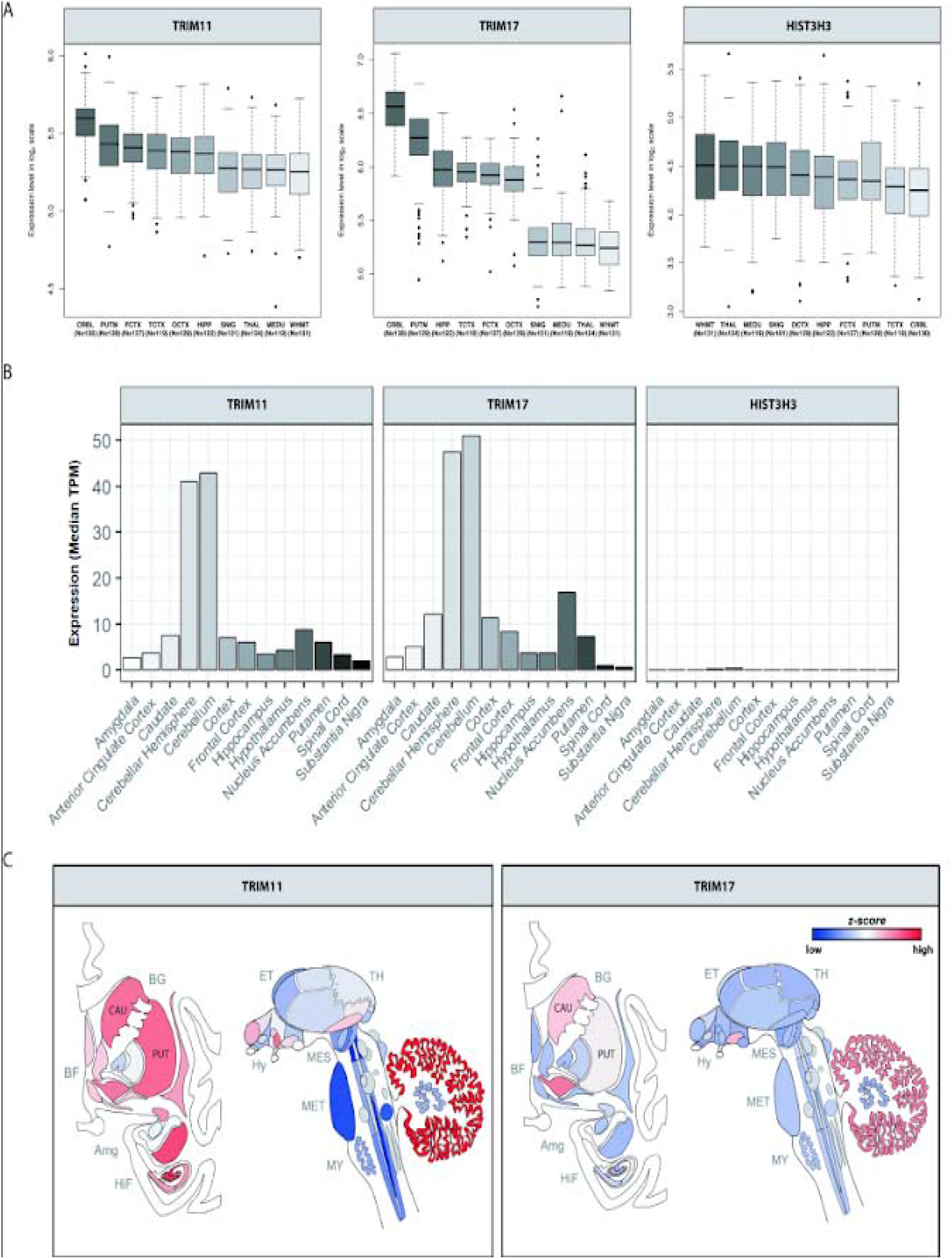
TRIM11, TRIM17 and HIST3H3 brain expression in: **A.** BRAINEAC database – CRBL = Cerebellum, PUTM = Putamen, HIPP = Hippocampus, TCTX = Temporal cortex, OCTX = Occipital cortex, FCTX = Frontal cortex, THAL = Thalamus, SNIG = Substantia nigra, MEDU = Medulla, WHMT = White matter; **B.** GTEx database; **C.** Allen Atlas database (Caucasian subjects) – Amg = Amygdala, BF = Basal Forebrain, BG = Basal Ganglia, CAU = Caudate, ET = Epithalamus, HiF = Hippocampal Formation, Hy = Hypothalamus, MES = Mesencephalon, MET = Metencephalon, MY = Myelencephalon, PUT = Putamen, TH = Thalamus. Image credit: Allen Institute.

We explored the cellular specificity of TRIM11, TRIM17 and HIST3H3 expression in human brain using data provided by the Brain RNA-seq database. This demonstrated higher neuronal expression (n = 1) of TRIM11 (0.38 FPKM) compared to both TRIM17 (0.12 FPKM) and HIST3H3 (0.10 FPKM), which was expressed at the lower limit of detection. In comparison to its neuronal expression, TRIM11 expression in glial cell types was lower (mature astrocytes 0.14±0.02 FPKM, n = 12; microglia 0.12±0.02 FPKM, n = 3; oligodendrocytes 0.11±0.01 FPKM, n = 3). We also explored cell-specific expression of the TRIM11 and TRIM17 mouse orthologues across the brain using single cell RNA-seq data (http://dropviz.org/) generated from mouse brain tissue. This data suggested that expression of TRIM11 was highest in the spiny projection neurons (SPNs) of the striatum, with high expression in SPNs of both the “direct” and “indirect” pathways. In contrast, TRIM17 expression was generally lower with the highest expression detected in the neurons of the substantia nigra.

Our colocalisation analysis did not reveal any significant associations between our GWAS signals and eQTL data in BRAINEAC and GTEx databases in all brain regions. However, of note, expression analyses using GTEx data revealed that several SNPs from the chromosome 1q42.13 locus reaching genome wide significance in our GWAS were significant eQTLs for TRIM11 and TRIM17 in skin and thyroid tissues when analysed individually.

## Discussion

To our knowledge, this is the first GWAS of clinical phenotype in PSP. We show that variation at the chromosome 1q42.13 locus determines clinical phenotype in PSP with a very strong effect size (odds ratio 5.5). The validity of our GWAS results is increased by the fact that similar sized association signals and minor allele frequencies were observed in two independent cohorts with a genome-wide significant association achieved when the two cohorts were combined. Furthermore, when considering the subset of MDS criteria phenotyped cases from the original PSP-case control GWAS, RS and non-RS group MAFs of directly genotyped SNPs in high LD with our lead SNP are supportive of our findings. We are also reassured by our PD genotyping data which revealed that our GWAS signal would have been attenuated if PD cases had been inadvertently included in our clinically diagnosed non-RS group. However, this is unlikely as we only included PSP-P and PAGF cases that fulfilled probable PSP MDS criteria for these phenotypes. We suspect that none of our significant SNPs reached genome wide significance in the original PSP case-control GWAS due to the fact that pathologically diagnosed PSP cases would have contained a mixture of RS and non-RS cases. When considering our data, we can infer that the combined MAFs of RS and non-RS cases would have resulted in overall PSP group MAFs that were similar to that of healthy controls. The validity of our NeuroChip genotyping and imputation were confirmed by the additional genotyping we carried out to span the chromosome 1q42.13 locus. The validity of our independent cohorts are suggested by the following: 1) In both cohorts, a majority of cases with an initial non-RS phenotype had a final clinical diagnosis of RS, as previously shown by other groups (28); 2) Our cohorts had similar MAFs for risk variants identified in the PSP case-control GWAS (15); 3) There was 100% concordance between clinical and pathological diagnoses in our clinical cohort for the subset of patients that had undergone post-mortem.

MAGMA analysis confirmed signals in *TRIM11, TRIM17* and *HIST3H3* genes. There was evidence for differential regional brain expression of TRIM11 and TRIM17. Both human and mouse RNA-seq data revealed high levels of neuronal TRIM11 expression. In addition, it is likely that our GWAS was significantly underpowered to detect signals in our eQTL colocalisation analyses.

It is important to note that SNPs within this genomic locus are in high LD, as evidenced by the spread of genome wide significant SNPs identified in our GWAS (Figure 2). Therefore, it is challenging to know which gene is driving our association signal. However, the localisation of the lead SNP in our dataset and the gene expression profiles described above suggest that *TRIM11* is the most likely candidate gene at the chromosome 1q42.13 locus. The eQTL profile of our significant SNPs in GTEx, when analysed individually, was particularly interesting. The strong association between several SNPs in high LD with rs564309 and decreasing TRIM11 and TRIM17 expression in non-brain tissues highlight the concept of tissue/region/cell specific expression of transcripts potentially being determined by disease state and at specific time points in development or ageing (29). However, our data does not exclude potentially important functional roles for the other transcripts within this locus.

The major limitation of our study is the fact that our cohort size was relatively small compared to case-control GWAS in PSP (15) and other neurodegenerative diseases (30, 31). Further replication of our findings in larger cohorts is desirable, including other non-RS phenotypes such as PSP-F, to confirm the role of TRIM11 and identify genetic determinants of clinical phenotype in PSP at other loci. Furthermore, our lead SNP is an intronic variant that does not pass the FDR threshold for being a brain eQTL at the genes in our locus of interest and the only coding variants that it is in LD with are from a gene (*OBSCN*) that is unlikely to be of biological relevance to PSP pathology. This is a common dilemma as a majority of risk variants identified in GWAS over the past two decades are not associated with coding changes in expressed proteins (32). Furthermore, disease associated intronic SNPs can regulate the expression of more distant genes. When referring to BRAINEAC and GTEx, rs564309 was found not to be a brain eQTL at distant genes outside of our region of interest. Indeed, functional impacts of intronic variants may arise in modes other than gene expression, including via splicing and methylation patterns of targeted transcripts and proteins. It remains a challenge to understand the functional consequences of non-coding genetic variation linked to phenotype, and so functional studies are vital. However, gene expression studies in post-mortem disease tissue can be challenging to interpret because of the confounding effects of changes on cell populations (33).

TRIM proteins are biologically plausible candidates as determinants of clinical phenotype in PSP and promising targets for follow up functional studies. The TRIM family of proteins, most of which have E3 ubiquitin ligase activities, have various functions in cellular processes including intracellular signalling, development, apoptosis, protein quality control, autophagy and carcinogenesis (34). A recent study has shown that TRIM11 has a critical role in the clearance of misfolded proteins via the ubiquitin proteasome system (UPS), in this case pathogenic fragments of both Ataxin-1 (Atxn1 82q) and Huntingtin protein (Httex1p 97QP) (35). Other groups have shown that lysine residues of tau are targets for polyubiquitination which induces proteolytic degradation of tau via the UPS (36). Furthermore, tau accumulation has been associated with decreased proteasome activity in mouse tauopathy models, suggesting a feedback loop between impaired protein degradation, aggravated by a protein aggregate based impairment of proteostasis (37). These findings coincide with previous studies that have identified the UPS as a potential drug target in the treatment of neurodegenerative conditions (38, 39). Overexpression of TRIM17, partly controlled by glycogen synthase kinase 3 pathways, has been shown to initiate neuronal apoptosis in cell models (40). This was later shown to be mediated by increased degradation of the anti-apoptotic protein, myeloid cell leukaemia 1 (Mcl-1), via the UPS (41).

Based on our data, we hypothesise that common variation at the chromosome 1q42.13 locus modifies the function of TRIM11 to varying degrees in specific brain regions. In the more slowly progressing non-RS syndromes, an increase in TRIM11 function may lead to increased degradation of toxic tau species *via* the UPS, therefore protecting against tau pathology. Conversely, a decrease in protein function in the brainstem is more likely to promote rapid accumulation of tau aggregates, manifesting as the malignant RS phenotype of PSP.

In summary, the results of this study suggest that common variation at the *TRIM11* locus may be a genetic modifier of clinical phenotype in PSP. Our findings add further evidence for the UPS playing a key role in tau pathology and therefore representing a potential target for disease modifying therapies. Further GWAS with larger cohorts to confirm our findings and identify other genetic signals, screening of whole genome/exome sequencing data for rare variants in *TRIM11* and follow up functional studies at this locus are a priority.

## Acknowledgements

We would like to acknowledge the contribution of all PROSPECT-UK investigators for their role in acquiring patient data that was vital to this study. PROSPECT-UK is primarily supported by the PSP Association.

EJ and JW are supported by the PSP Association. MMXT is supported by Parkinson’s UK. MS is supported by Cytox Limited and Innovate UK. AP is supported by the University College London Hospitals Biomedical Research Centre (BRC). RF is supported by the Alzheimer’s Society. KYM is supported by the Reta Lila Weston medical trust. DZ, RdS and MR are supported by the Medical Research Council. RHR is supported through the award of a Leonard Wolfson Doctoral Training Fellowship in Neurodegeneration. HRM is supported by the PSP Association and CBD Solutions. All other authors did not declare any funding sources that directly contributed to this study.

## Author Contributions

Study concept and design - EJ and HRM; Data acquisition and analysis – EJ, JW, MMXT, MS, AP, RF, KYM, DZ, RHR, RdS, MJG, GR, UM, SAS, SMG, TR and JLH; Drafting and critical analysis of manuscript - EJ, AJL, TTW, JH, TR, GUH, JLH, MR and HRM.

